# Alternate models of acute dyslipidemia reveal divergent pathways upon atherosclerosis initiation

**DOI:** 10.1101/2020.09.02.279596

**Authors:** Sanna Hellberg, Vladimir S. Shavva, Jari Metso, Guangchun Chen, Quan-Zhen Li, Valentina Manfè, Peder S. Olofsson, Matti Jauhiainen, Christoph J. Binder, Stephen G. Malin

## Abstract

Atherosclerosis is thought to be initiated by the sub-intimal retention of apolipoprotein-B containing lipoproteins within susceptible sites of the vasculature. Understanding this initiation is not possible with current legacy mouse models of atherosclerosis. We created two mouse strains of inducible hypercholesterolemia based on conditional loss of apolipoprotein E or through inducible expression of a gain-of-function proprotein convertase subtilisin/kexin type 9 D374Y mutation. Both strains rapidly broke plasma-lipid homeostasis and converted to a state of atherogenic dyslipidemia, resulting in overt aortic-accumulation of lipoproteins within 10 days. RNA-sequencing revealed that the vascular response is completely dependent on the route taken to dyslipidemia, which nevertheless implicates known pathogenic pathways in the aetiology of atherosclerosis, and further implicates APOE as an inhibitor of inflammation. As atherosclerosis develops, a convergence of common mechanistic processes emerge in both strains with significant involvement of the immune system, and targeting of CD8 T cells can regulate a conserved aortic response. Our results define atherosclerosis initiation as highly heterogeneous process and identify multiple potential therapeutic targets that may influence disease onset.

## Introduction

Atherosclerosis is a major underlying cause of coronary heart disease and stroke ^1^. There is overwhelming evidence that atherogenesis is instigated by increased levels of circulating apolipoprotein B containing lipoproteins (apoB-lipoproteins), most notably low-density lipoprotein (LDL) ^2^. In the current paradigm, apoB-lipoproteins are retained in the arterial intima at sites where the endothelium is dysfunctional. This endothelial dysfunction can be caused by low shear stress or mechanical injury, and it is characterized by increased permeability and altered expression of adhesion molecules ^3^. Retained lipoproteins can be modified and aggregated, resulting in the generation of epitopes that are recognised as dangerous by cells present in blood vessels, and which can subsequently trigger an immune response ^4^. This results in further recruitment of inflammatory cells from the bloodstream, mainly monocytes but also dendritic cells (DC) and lymphocytes ^5^. Arterial resident cells, particularly smooth muscle cells, can also be modified during atherogenesis ^6^.

Macroscopically, the first manifestation of atherosclerosis is the formation of fatty streaks, which largely consist of monocyte-derived macrophages that have become engorged with lipid due to excess uptake and storage of modified lipoproteins ^7^. The *in vivo* responses in atherosclerosis-susceptible arterial sites to raised blood cholesterol levels prior to fatty streak formation has remained experimentally inaccessible and largely unexplored.

Wild-type mice have low LDL levels and are resistant to atherosclerosis unless subjected to specialized diets over greatly extended periods of time ^8^. Current mouse models for atherosclerosis are therefore based on congenital defects in lipoprotein uptake mechanisms, most commonly apolipoprotein E deficient (*Apoe*^−/−^) or LDL receptor deficient (*Ldlr*^−/−^) strains that present markedly increased plasma apoB-lipoprotein concentrations ^9–11^. *Apoe*^−/−^ mice develop atherosclerotic lesions even on chow diet, with fatty streaks visible from 8 weeks of age ^12^, whereas *Ldlr*^−/−^ mice are considered to require a high-cholesterol diet in order to become atherosclerotic within reasonable time frames ^13^. The extent that observations from one individual strain are applicable to atherosclerosis in general is a matter of debate ^14^. Nevertheless, mouse models have proven exceptionally useful in revealing the underlying mechanisms behind atherosclerotic plaque formation ^15^.

Due to the chronic nature of the most commonly used mouse models, the mechanisms initiating plaque formation have remained obscure. For example, exogenously supplied apoB-lipoproteins are retained in the rabbit aorta two hours after injection ^16^, and *Apoe*^−/−^ mice at weaning already display lipid accumulation in atherosclerosis prone regions of the aorta ^17^. This would indicate that the first step in atherogenesis, namely retention of apoB-lipoproteins within the arterial intima, occurs rapidly following increased plasma cholesterol levels. However, this process cannot be interrogated with legacy dyslipidemic strains, as both *Apoe*^−/−^ and *Ldlr*^−/−^ mice present constantly elevated and allostatic plasma apoB-lipoprotein levels. We rationalised that acutely transitioning to a state of dyslipidemia would provide a window in which to observe the microscopic and molecular transitions that typify atherosclerosis initiation. To achieve this, we utilized two different mouse models of acute dyslipidemia, an inducible *Apoe* knockout ^18^ and a novel model with inducible expression of a human proprotein convertase subtilisin/kexin type 9 (hPCSK9) D374Y gain-of-function variant ^19–21^. Herein, we challenge the hypothesis that a singular pathway demarcates the onset of atherosclerosis. We provide evidence that the aortic response to a dyslipidemic insult is dependent on the defect in lipoprotein uptake that breaks homeostasis of plasma cholesterol metabolism. However, common pathways become apparent as atherogenesis progresses and these can be modified by targeting immune components. Taken together, these findings reveal an intricate interplay between the vasculature and the immune system upon atherosclerosis initiation.

## Results

### Parallel mouse modelling to engender a state of dyslipidemia

We have previously shown that inducible loss of APOE in the adult mouse leads to acute hypercholesterolemia ^18^. We created a complementary mouse model based on the conditional expression of a human proprotein convertase subtilisin/kexin type 9 gain-of-function variant (hPCSK9 D374Y) inserted into the ROSA26 locus. The tamoxifen inducible and ubiquitously expressed *Cre* line *ROSA26*^*CreERt2/ CreERt2*^ was additionally crossed with this hPCSK9 D374Y strain. Hence both conditional loss of *Apoe* (*Apoe*^*fl/−*^ *ROSA26*^*CreERt2/+*^ herein referred to as APOE cKO, with *Apoe*^*fl/+*^*ROSA26*^*CreERt2/+*^ as littermate controls) and inducible expression of hPCSK9 D374Y (*ROSA26*^*CreERt2/hPCSK9D374Y*^ herein referred to as D374Y, with *ROSA26*^*CreERt2/+*^ as littermate controls) can be achieved by tamoxifen administration (**Fig1a**). We determined hPCSK9 levels by enzyme-linked immunosorbent assay (ELISA) before tamoxifen administration in the 10-week old D374Y and littermate control mice (*ROSA26^CreERt2/ CreERt2^* or *ROSA26*^*CreERt2/+*^ herein referred to as littermate controls) and both had similar levels of hPCSK9 protein in the plasma (199 vs. 191 ng/ml). This increased 11-fold (2325 vs. 213 ng/ml) at three days post-tamoxifen, (**FigS1a**). The hPCSK9 plasma levels were therefore at levels physiologically relevant in humans ^22^. The plasma cholesterol levels followed the hPCSK9 expression and reached maximum already at 3 days after tamoxifen dosing (**FigS1b**).

**Figure 1.**
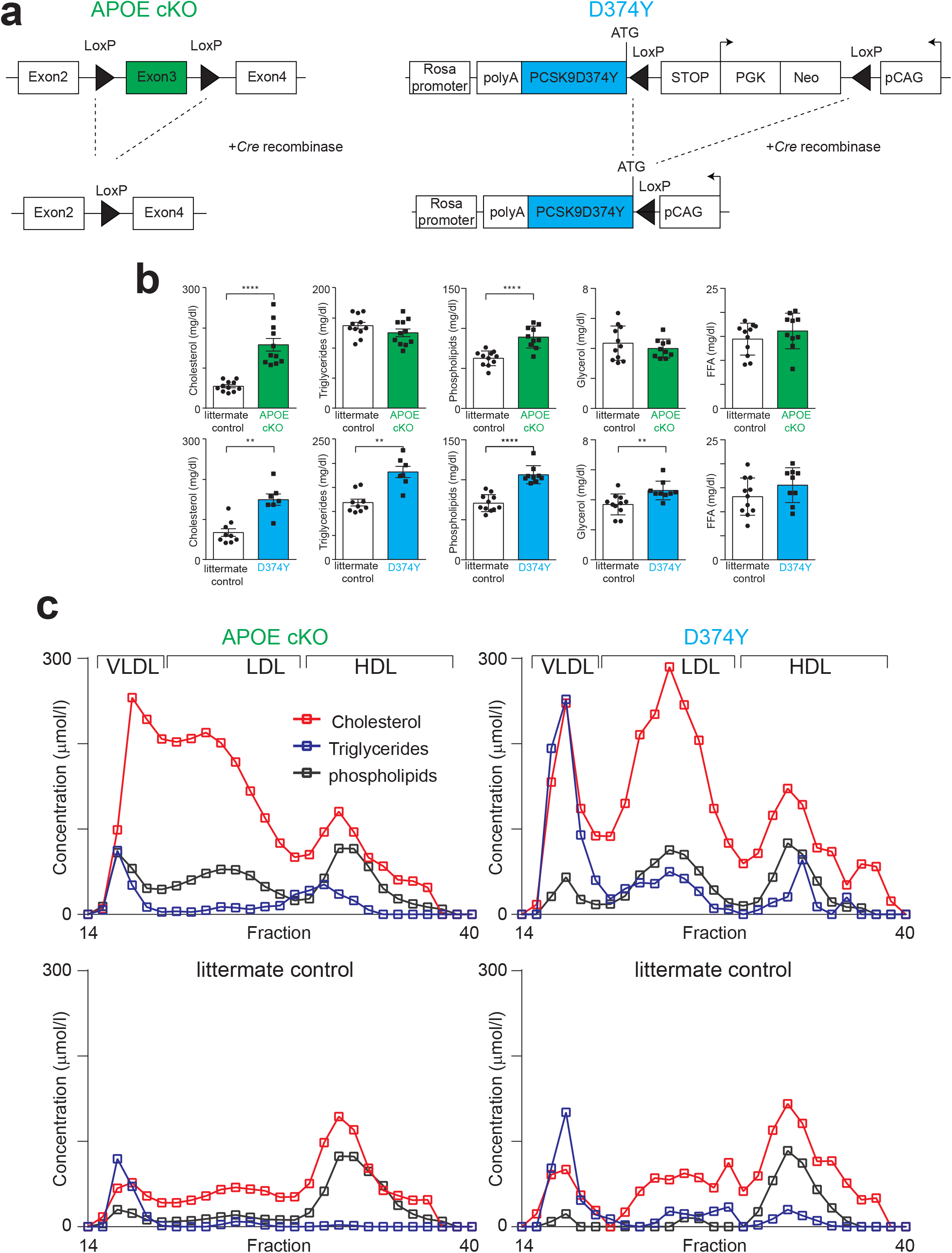
Generation and characterization of mouse models for inducible dyslipidemia. **A.** Schematic diagram of the inducible Apolipoprotein E knockout (APOE cKO) and human proprotein convertase subtilisin/kexin type 9 (hPCSK9) D374Y variant (D374Y) alleles. In the APOE cKO strain, Exon 3 of *Apoe* is flanked by *loxP* sites and is deleted upon *Cre* recombinase activity from *ROSA26^CreERT2/+^*. The second allele is germline deleted for *Apoe,* producing *Apoe*^*fl/−*^ ROSA26^*CreERT2/+*^ mice, with *Apoe*^*fl/+*^ ROSA26^*CreERT2/+*^ used as littermate controls. For D374Y, the coding sequence of hPCSK9 containing the mutation D374Y was inserted into the ubiquitously expressed *Rosa26* locus that additionally contains the CAG promoter, together with a *loxP* flanked stop cassette. This allele was combined with *ROSA26*^*CreERT2/+*^ to produce *ROSA26^*CreERT2/hPCSK9D374Y*^*, with *ROSA26^*CreERT2/CreERT2*^* as littermate controls. In D374Y mice, *Cre* expression deletes the stop cassette and expression of hPCSK9 D374Y is initiated. Neo, neomycin; PGK, phosphoglycerate kinase. **B.** Plasma lipid levels (mg/dl) 10 days after tamoxifen administration in APOE cKO (green bars) and D374Y (blue bars) strains together with respective littermate controls in male and female mice fed normal chow. Differences between control and experimental mice calculated by Student’s t test. APOE cKO and littermate controls (n = 10-11). D374Y and littermate control (n = 7-11). All univariate scatter plots are ± SEM. **p < 0.01 and ****p < 0.0001 compared to littermate control mice. **C.** Plasma lipoprotein fractionation profiles (μmol/l) at 10 days after tamoxifen dosing. All curves calculated as an average of two separately run plasma pools from male and female mice (plasma from 4-6 mice in each pool).

We next compared the plasma lipids in these two strains at day 10 after tamoxifen administration, with mice maintained on a normal chow diet (**Fig1b**). Cholesterol levels were increased in both APOE cKO (158 mg/dl vs 55mg/dl in littermate controls) and D374Y (150 mg/dl vs 67 in littermate controls), as were phospholipid levels (89 vs 63 mg/dl in APOE cKO and controls, and 106 vs 71 mg/dl in D374Y and controls). Noticeably, the D374Y strain displayed increased plasma levels of triglycerides and glycerol, which was not observed in the APOE cKO mice (**Fig1b**). Circulating free fatty acid levels were comparable in all four genotypes, and the levels of all measured lipid species in the two control groups were similar to each other. Finally, we determined which lipoprotein species were responsible for the observed increases in cholesterol and triglycerides through fast-performance liquid chromatography. In both the APOE cKO and D374Y strains, cholesterol was incorporated in the VLDL/chylomicron remnant fraction of lipoproteins, whereas in D374Y a more prominent accumulation in the LDL fraction was additionally present (**Fig1c)**. The increase in triglycerides present in D374Y mice were incorporated into the VLDL/chylomicron remnant fraction. In summary, acutely induced dyslipidemia in adult APOE cKO and D374Y mice reassemble those of the legacy *Apoe*^−/−^ and *Ldlr*^−/−^ strains ^9–11,23^. We have therefore created two strains that allow for either acute inducible hypercholesterolemia (APOE cKO) or mixed dyslipidemia (D374Y), in the adult mouse.

### Rapid retention of Apolipoprotein B at sites of atherosclerosis susceptibility following a transition to dyslipidemia

Retention of apoB-lipoproteins in the arterial intima is considered the starting point of atherosclerotic plaque development. We next addressed if we could visualize this commencement of atherogenic lipid retention, in aortas collected *ex vivo* from APOE cKO and D374Y at 10 days after tamoxifen treatment. To complement our inducible models, we also included aortas from adult *Apoe*−/− and *Ldlr*−/− mice aged 10 weeks for the confocal microscopy analysis. All the mice were maintained on a chow diet. Increased amounts of APOB was readily detectable in the atherosclerosis-prone lesser curvature, but not greater curvature, of aortic arch at 10 days post-tamoxifen in both inducible models but not littermate controls (**Fig2a).** Higher magnification images clearly demonstrated sub-intimal retention of APOB in close proximity to CD68+ cells (**Fig 2b**). We quantitated APOB retention along one millimetre of the atherosclerosis-prone lesser curvature in ascending aorta, and observed increased APOB retention in both inducible models (**Fig2c**). Noticeably, 10-week old *Apoe*−/− and *Ldlr*−/− strains presented even more retained APOB (**Fig 2c**), in agreement with the higher cholesterol levels and the sustained dyslipidemia (10 weeks) of these mice (**Fig2d**). CD68+ cells in D374Y but not APOE cKO mice additionally displayed increased cell size, indicative of activation **(Fig2e)**. In summary, these inducible mouse strains provide a stereotypical transition from a healthy aorta to the first initiation stage of atherosclerosis, as manifested by sub-intimal APOB retention, and this stage of atherosclerosis initiation is not amenable to experimentation in *Apoe*^−/−^ and *Ldlr*^−/−^ mice.

**Figure 2.**
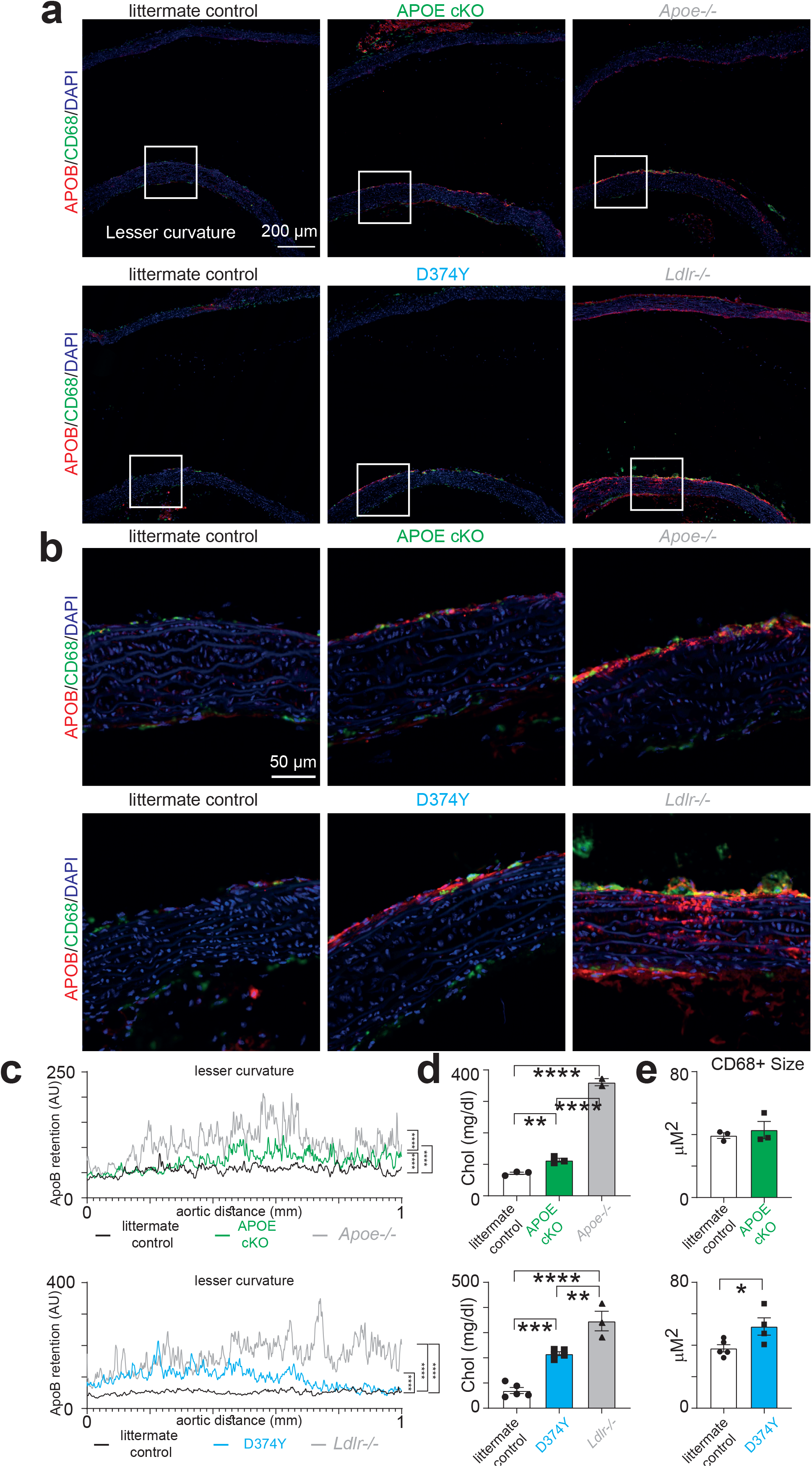
Rapid accumulation of apolipoprotein B in atherosclerosis-susceptible arterial sites in response to dyslipidemia. **A**. Confocal microscopy of the ascending aorta in APOE cKO, D374Y and their respective littermate controls at 10 days after tamoxifen dosing, as well as 10-week old apolipoprotein E deficient and low-density lipoprotein receptor deficient mice. All mice were on chow diet and mixed genders. Immunofluorescence staining for apolipoprotein B (APOB) (red) and CD68 (green) is shown at 10x magnification. Nuclei are visualized by 4′,6-diamidino-2-phenylindole (DAPI) (blue). **B.** Close-up of regions outlined with dashed lines from A. **C.** Quantitation of APOB staining in one millimetre stretch of the lesser curvature intima in ascending aorta (AU= arbitrary units). **D**. Plasma total cholesterol levels of the mice analysed. **E.** Size measurements (μm^2^) of CD68+ cells located in the lesser curvature of the aorta.Statistical data (B–E) are shown as mean value with SEM and analyzed by repeated measures one-way ANOVA (C), one-way ANOVA (D) or Student’s t-test (E); *p < 0.05, **p < 0.01, ***p < 0.001, ****p < 0.0001 compared to control mice.

### Molecular profiling of atherosclerosis initiation

The creation of inducible systems to trigger intimal APOB retention provided us with the opportunity to discover the vasculature’s response to atherosclerosis onset. Therefore, we next performed a series of mRNA-seq experiments on thoracic aortas to determine the gene expression changes upon atherosclerosis initiation. We first compared APOE cKO with littermate controls at the very early time point of 7 days following tamoxifen administration (**Fig3a)**. 27 genes were differentially regulated (individual P-value of < 0.002, false discovery rate (FDR) < 0.1 and normalized gene expression (transcripts per million reads mapped, TPM) > 5 in either all control or experimental samples), including strong downregulation of *Apoe* as expected (**Fig3b)**. The most strongly regulated genes could be classified into those regulating angiogenesis and vascular function (*Serpine1, Rnh1, Kdr, Btg, Creb3l2, Chchd2*), metabolism (*Apoc1*, *Alas2, Elovl3*) and T cell infiltration (*Cd8a,* **Fig3c**). Hence, even this very short time period of hypercholesterolemia was sufficient to change the transcriptional make-up of the aorta.

**Figure 3.**
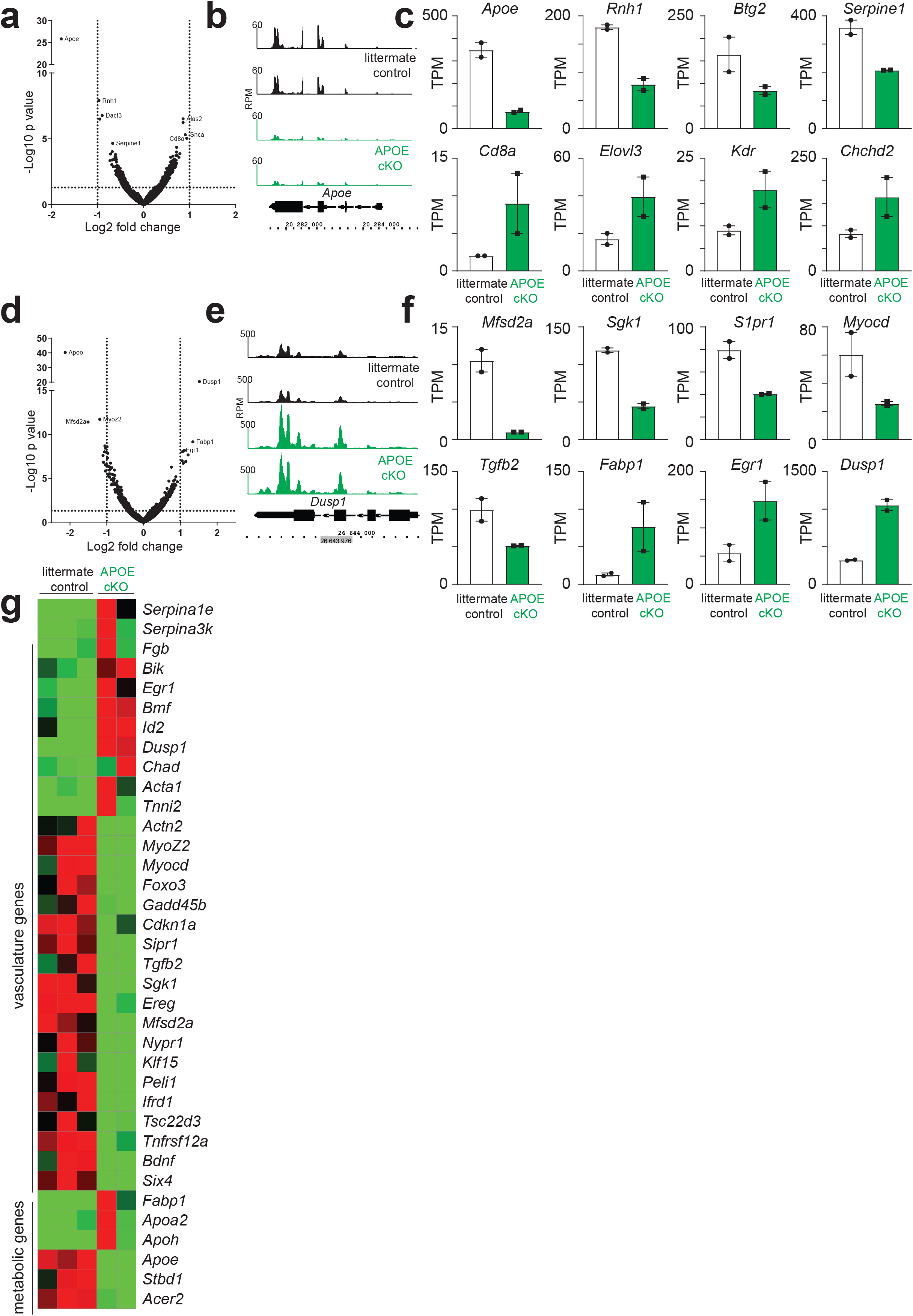
Aortic gene expression changes following acute dyslipidemia in APOE cKO mice. **A.** Volcano plot of aortic gene expression differences in female APOE cKO mice compared to littermate controls at day 7 after tamoxifen dosing, based on two RNA-seq experiments for each genotype. Selected genes with an adjusted P-value of < 0.1, and a TPM value of > 5 (in one aorta samples) are indicated. **B**. Mapped reads (RPM) of the *Apoe* locus from the APOE cKO (green) and littermate controls (black). **C**. Expression of selected significantly activated and repressed genes from APOE cKO (green) and littermate control female aortas. The mRNA expression of the indicated genes is shown as mean expression value (TPM) with SEM, based on two different RNA-seq experiments for each genotype. **D.** Volcano plot of gene expression day 10 after induction of dyslipidemia, **E**. Mapped reads (RPM) of the *Dusp1* locus. **F**. Expression of selected differentially regulated genes from female APOE cKO (green) and littermate controls aortas based on three different RNA-seq experiments from littermate controls and two from APOE cKO. **G.** Heatmap of selected genes ordered by function in APOE cKO and their respective littermate controls. Red indicates higher expression and green lower expression normalised for each individual gene.

At 10 days after tamoxifen administration, more striking patterns of gene expression were observed in APOE cKO mice, with 113 genes differentially regulated (P-value < 0.001, FDR < 0.1 and TPM > 5, **Fig3d,e**). Three separate groupings could be discerned, which included metabolic genes of the apolipoprotein family (*Apoa2; Apoh; Apoe*). However, most noticeable was the dysregulation of genes involved in platelet biology and coagulation (*Serpina1e; Fgb; Serpina3k*) and multiple established genes with functions in vascular identity and function (*Tnni2; Actn2; Acta1; Cdkn1a; Chad; Dusp1; Foxo3; Ereg; Bdnf; Sgk1; Gadd45b; S1pr1; Tgfb2* **Fig3f,g)**.

The D374Y strain showed a markedly different gene expression response following induction of dyslipidemia, with no overlap of gene expression changes in the aorta of D374Y mice compared to controls with those of the APOE cKO analysis, at day 10 after tamoxifen administration. The majority of transcriptional changes resulted in upregulation of genes in the D374Y strain (86 out of 89, P < 0.0001, FDR < 0.15, TPM > 5) (**Fig4a**). These included multiple metabolic factors, especially those central to the fatty-acyl-CoA and triglyceride biosynthetic process (*Acsm3, Acly, Acsl1, Ehhadh, Dlat, Dgat1, Nudt7, Acsl5, Lpl, Scd1, Dbi*) and a number of genes with known function in cardiovascular disease (*Lamb1, Adipoq, Rasd1, C3*). However, most striking was the upregulation of genes central to immune responses in myeloid cells. This included H-2 class II histocompatibility antigen genes (*H2-Aa, H2-Eb1, H2-Ab1*), cell surface receptors that recognize native and modified lipids (*Cd1d1, Cd36*) and multiple mediators of mast cells and myeloid activation (*Orm1, Mpeg1, Ptger3, Mafb, Mgst1, Iigp1, Pld4, Cma1* (**Fig4b,c**). These differences are summarized as heat maps in **Fig4d.** Taken together, these results indicate that atherosclerosis initiation is characterized by two distinct gene expression programs in the aorta dependent on which route to dyslipidemia is taken; with hypercholesterolemia in the APOE cKO modulating vasculature and platelet-coagulation pathways, and the mixed dyslipidemia of the D374Y strain characterized by upregulation of metabolic and immune system genes.

**Figure 4.**
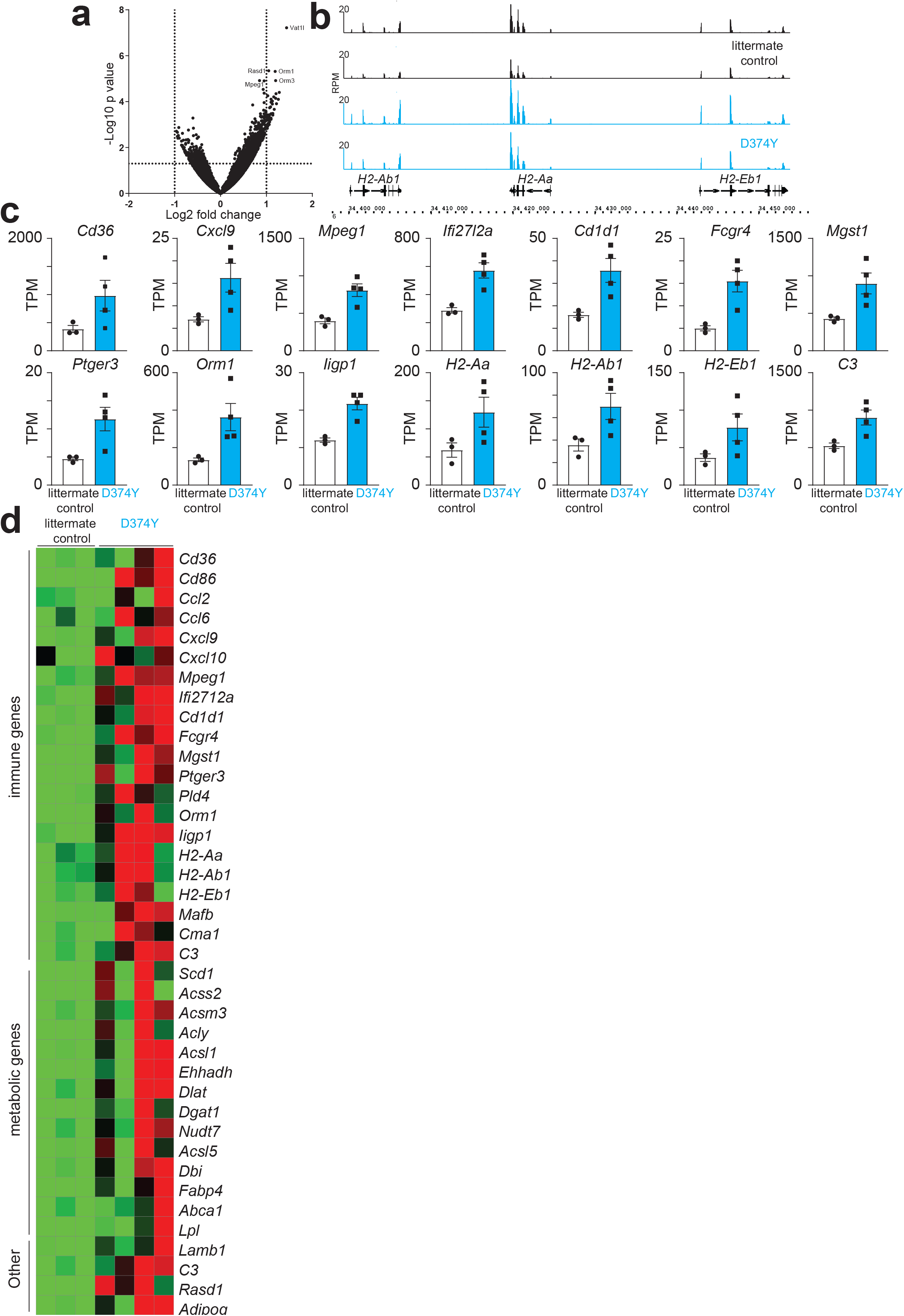
Divergent patterns of aortic gene expression in response to dyslipidemia. **A.** Volcano plot of gene expression differences in D374Y (4 RNA-seq experiments) female mice compared to littermate controls (3 RNA-seq experiments) at day 10 after tamoxifen dosing. Selected genes with an adjusted P-value of < 0.15, and transcripts per million reads mapped (TPM) value of > 5 (in one aorta samples) are indicated. **B**. Mapped reads (RPM) across part of the *MHCII* locus from D374Y (blue) and littermate controls (black). **C**. Expression of selected activated and repressed genes from D374Y (blue) and littermate controls aortas. The mRNA expression of the indicated genes is shown as mean expression value (TPM) with SEM. Each dot corresponds to one RNA-seq experiment from female mice. **D**. Heatmap showing the relative expression of genes, clustered into immune, or metabolic or vascular function, in D374Y and their respective littermate controls. Red indicates higher expression and green lower expression normalised for each individual gene.

### Heterogeneity in immune populations and the plasma secretome dependent on the route to dyslipidemia

Loss of APOE is known to have multiple effects on the immune system, including the formation of splenic germinal centers (GC) ^24,25^ resulting in increased atherosclerosis ^18,26^. Therefore, we wished to determine whether there is a conserved immune response to dyslipidemia itself, by characterizing the immune compartment of our two mouse strains by flow cytometry of spleen cells. Analysis of a large cohort of mice at 10 days following tamoxifen administration revealed an approximate 2-fold increase in GC formation in the APOE cKO as previously reported ^18^, whereas no significant increase in GC cell numbers could be detected in the D374Y strain relative to controls (**Fig5a,b)**, indicating that hypercholesterolemia alone is not sufficient to induce the formation of GCs at this time point and could rather be due to loss of APOE. The absolute cell numbers of the splenic B1 and B2 cell populations were not affected in either APOE cKO or D374Y mice. Similarly, the major CD4+ T cell populations and CD8+ cells were unaffected in either strain at day 10 following tamoxifen administration (**FigS2**). In contrast, the absolute number of DC populations showed a reciprocal pattern, with cDC1 cell numbers increased approximately one third in APOE cKO mice and decreased approximately 40% in the D374Y strain, relative to their specific littermate controls (**Fig5c,d)**. There was no changes in the absolute cell numbers of the cDC2 population. Expression of major histocompatibility complex-II (MHCII) was also unchanged on DCs in all genotypes.

**Figure 5.**
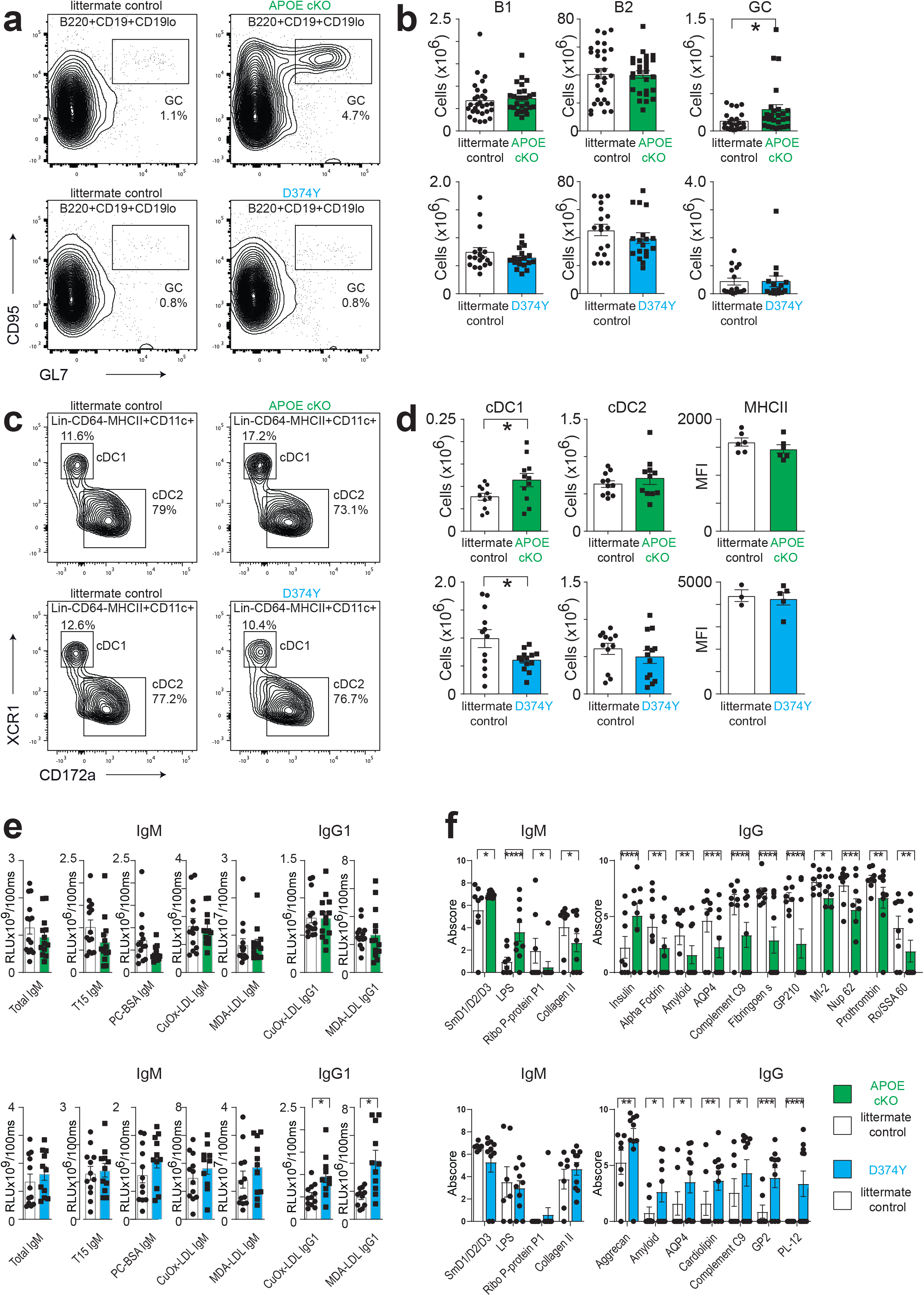
Immune responses to acute dyslipidemia. **A**. Flow cytometric analysis of germinal center (GC) B cells from APOE cKO, D374Y and their respective littermate controls at day 10 following tamoxifen administration. Percentage of cells within quadrants are indicated. **B**. Absolute cell numbers of B1, B2 and GC B cells from mice of the indicated genotypes. **C**. Representative plots of cDC1 and cDC2 cell populations. **D**. Cell numbers of dendritic cell populations of indicated genotypes and expression levels (MFI) of MHCII on gated CD11c+CD11b-CD19- dendritic cells. **E**. Plasma levels of total IgM, IgM against T15 idiotype and phosphorylcholine (PC-BSA), and IgM and IgG1 against Cu2+ oxidized (CuOx-LDL) and malondialdehyde modified low-density lipoprotein (MDA-LDL) in mice of the indicated genotypes at day 10 after tamoxifen dosing. Each dot corresponds to one mouse. **F.** Autoantigen arrays. IgM and IgG reactivity against selected autoantigens. Statistical data (B,D,E) are shown as mean value with SEM and analyzed by Student’s t-test; F two-way ANOVA *p < 0.05, **p < 0.01, ***p < 0.001, ****p < 0.0001 compared to control mice.

Using ELISA measurements, we next determined the extent of humoral responses against modified LDL upon the transition to acute dyslipidemia. Surprisingly, IgG1 reacting against both Copper (Cu2+) oxidized and malondialdehyde (MDA) modified LDL, was approximately 2-fold increased in D374Y mice relative to littermate controls (**Fig5e)**. This increase was not present in the APOE cKO, despite of their GC formation. Both strains showed no significant increase in IgM responses at this time point. We complemented the analysis of immune responses by measuring plasma IgM and IgG antibodies against 120 selected autoantigens by microarray. At day 10, the strains showed divergent autoantibody responses **(Fig5f**). IgM antibodies against SmD1/D2/D3 and LPS were increased in APOE cKO relative to controls, whereas reactivity against ribosomal phosphoprotein 1 and collagen II was decreased. In the D374Y strain, no significant changes in autoreactive IgM responses could be detected. IgG responses showed even more variability between the strains. APOE cKO displayed mostly reductions in autoantibody titers, with the exception of anti-insulin IgG. D374Y upregulated IgG reactivity to multiple self-antigens. Of note, this included amyloid and AQP4, both of which were conversely significantly down-regulated in APOE cKO mice.

Finally, we wished to test whether other plasma proteins components beyond antibodies were differentially regulated in the two strains. We analyzed 88 different plasma proteins by proximity extension assay. This revealed that secreted factors were mostly downregulated in the APOE cKO strain, including TGFB1, GDNF, IL17A and CXCL9. In contrast D374Y increased production of IL5 and GCG and decreased IL1A and PLXNA4 following induction of dyslipidemia (**Fig6a,b)**. Of the proteins measured, the secretion of only one factor was conserved between strains, which was a decrease in PDGFB (**Fig6b)**. In summary, the extent of conserved immune and humoral responses in the two strains appear to be limited to increased concentrations of plasma cholesterol-rich lipoproteins and decreased production of platelet derived growth factor subunit B.

**Figure 6.**
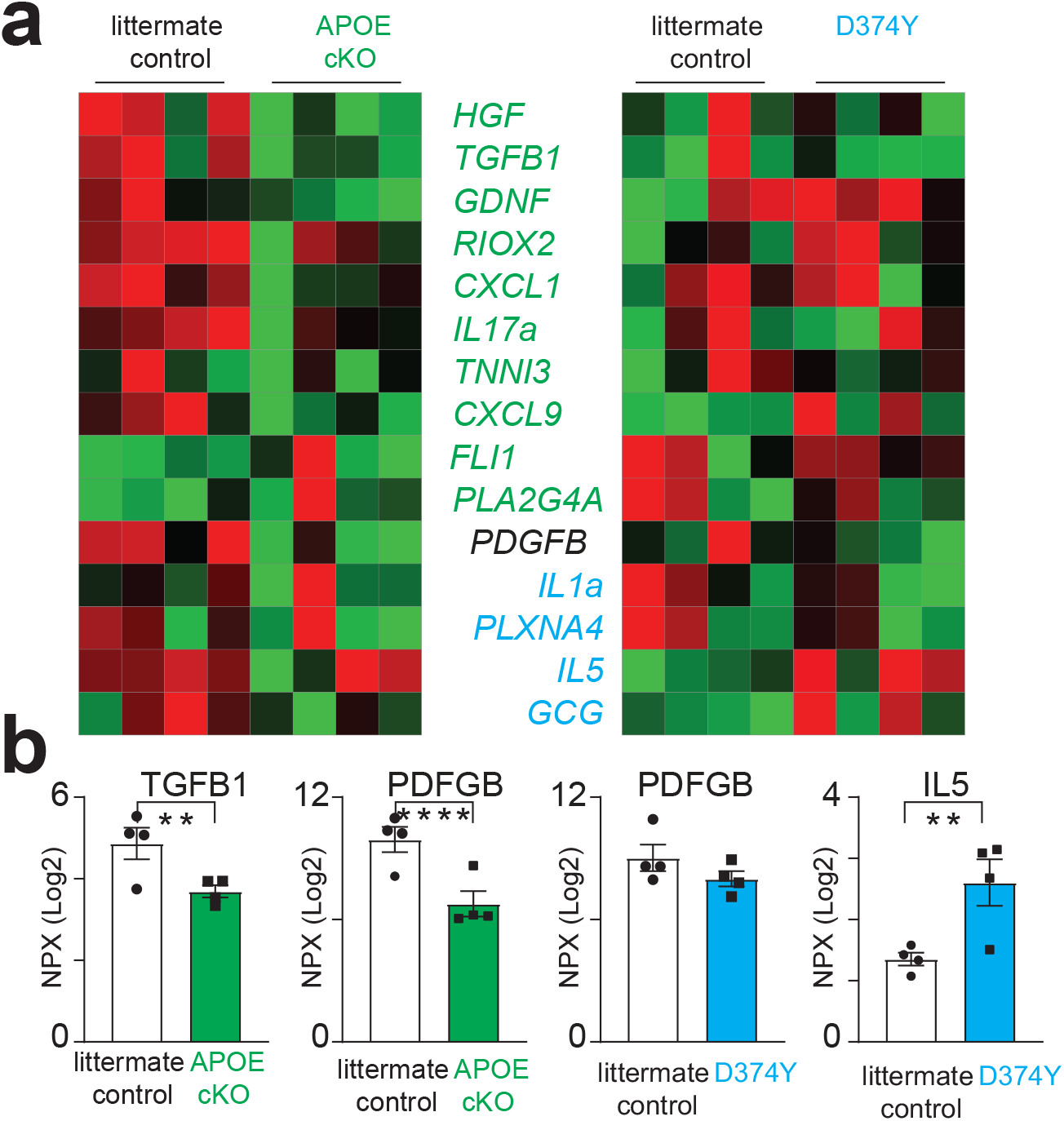
Analysis of plasma protein profiles in response to dyslipidemia. **A**. Heatmap of plasma proteins significantly differentially regulated in APOE cKO (written in green) and D374Y (written in blue) with their respective littermate controls at day 10 following tamoxifen administration as determined by proximity extension assay. Red indicates higher expression and green depicts lower expression levels normalized for each individual protein. **B**. Univariate scatter plots of selected proteins from the APOE cKO (green bars) and D374Y (blue bars) and their littermate controls. All plots are ± SEM and each dot corresponds to one mouse. Statistical analysis was performed with 2-way ANOVA. *p < 0.05, **p < 0.01, ****p < 0.0001 compared to control mice.

### Convergence of the aortic response during atherosclerosis progression

We next determined how the vasculature responds to sustained dyslipidemia by placing our two models on a high-fat diet (HFD) for 4 weeks immediately following induction of dyslipidemia. Comparison of aortic gene expression in APOE cKO with littermate controls identified 94 differentially regulated genes (21 down-regulated, 73 up-regulated, P < 0.0001, FDR < 0.1, TPM > 5 **Fig7a**). Only four genes, mostly metabolic regulators, were differentially expressed at both week 4 HFD and day 10 chow (*Apoe*; *Apoh*; *Fbp*1; *Slc10a1*). Although there was little overlap between specific genes at day 10 chow diet and week 4 HFD, genes involved in vasculature and platelet function as well as coagulation were again strongly enriched, including *Thbs1,Kng1, Pf4, Vsig4,* and *C3ar1* (**Fig7a)**. In strong contrast to the day 10 gene signature, the APOE cKO also upregulated multiple genes important for immune function, including *Itgb2, H2-Aa, Mrc1, Ctss, Fcer1g* and *Fcgr2b*. Analysis of the D374Y strain versus littermate controls after 4 weeks of HFD revealed greater changes in gene expression than compared to the day 10 chow fed mice, with 474 transcripts differentially regulated (172 down-regulated, 302 up-regulated P < 0.0001, FDR < 0.1, TPM > 5 **Fig7b)**. The D374Y strain again showed upregulation of immune genes, including *Cd24a, Cd38, Cd48, Cd53, Ly96* and *Clec7a*. Importantly, there was a conserved signature of immune activation genes in the aorta of D374Y mice at week 4 HFD with those at day 10, including *H2-Aa, Ctss* and *Mpeg1*, and the key metabolic regulators *Abca1, Dlat* and *Hsdl2* (**Fig7c**).

**Figure 7.**
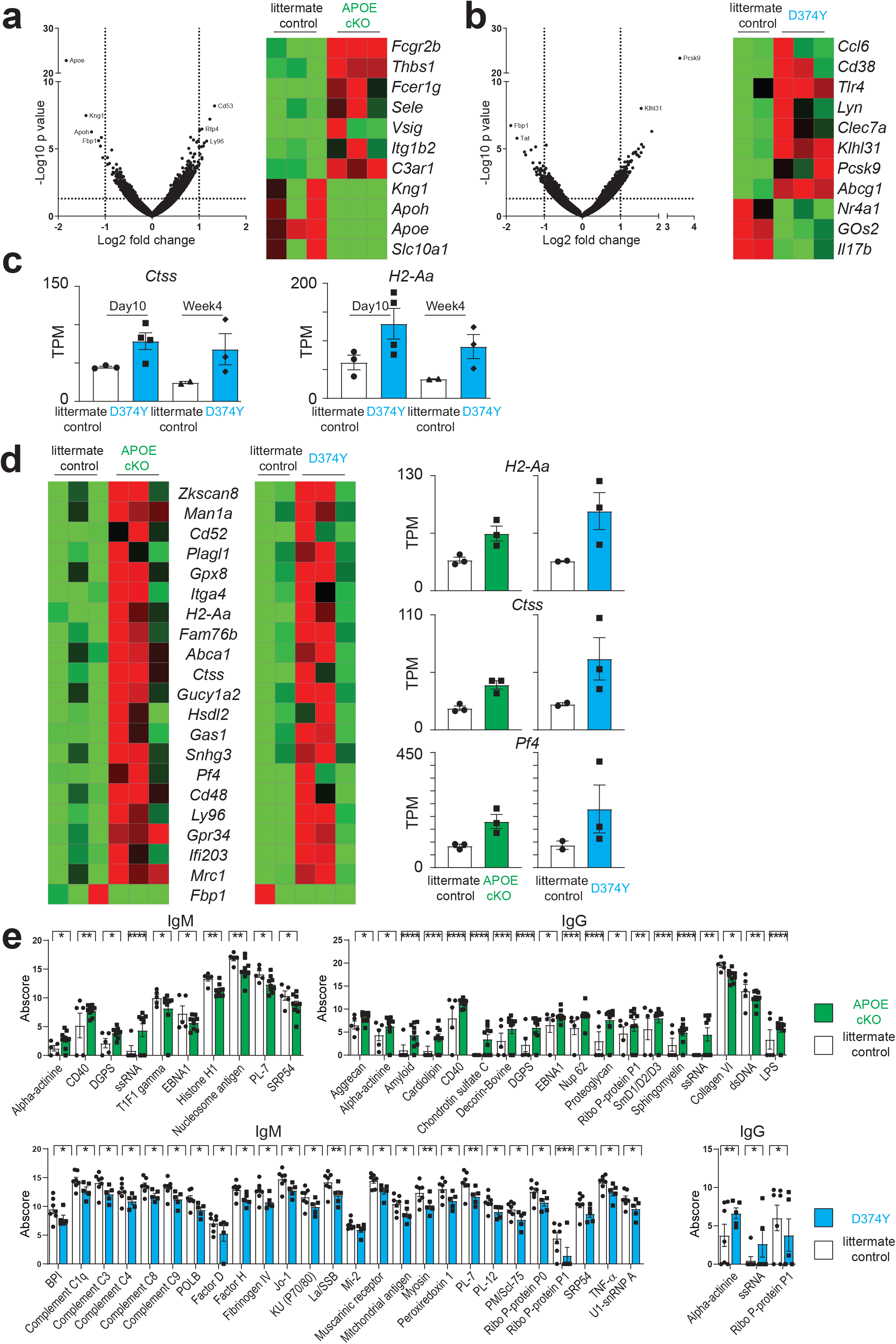
Convergence of gene expression patterns during atherosclerosis progression. **A,B** Left panel depicts volcano plot for gene expression differences in APOE cKO (3 RNA-seq experiments) mice compared to littermate controls (3 RNA-seq experiments), together with selected genes discussed in the text. Right panel shows D374Y (3 RNA-seq experiments) mice compared to littermate controls (2 RNA-seq experiments) and heatmap for selected differentially regulated genes. Both groups are female mice at 4 weeks following tamoxifen dosing and maintained on high fat diet. **C**. Selected genes significantly upregulated in D374Y and littermate controls both at day 10 and week 4 after induction of dyslipidemia. Each dot corresponds to a separate RNA-seq experiment from individual mouse aortas. **D**. Heatmap of conserved genes upregulated in both strains relative to controls in week 4 mice. Individual TPM for selected genes are depicted in the univariate scatter plots to the right. **E.** Autoantigen arrays. IgM and IgG reactivity against selected autoantigens for both strains and respective littermates. All plots are ± SEM and each dot corresponds to one mouse. Statistical analysis was performed with two-way ANOVA. *p < 0.05, **p < 0.01, ***p < 0.001, ****p < 0.0001 compared to control mice.

In strong contrast to the day 10 results, there was also an overlap in common gene expression signatures in both the APOE cKO and D374Y mice compared to respective littermate controls, with 21 genes being similarly dysregulated (**Fig7d)**. Hence over 20% of the genes differentially expressed in the APO cKO after 4 weeks of HFD were also similarly affected in the D374Y strain. These conserved dyslipidemia-responsive genes included the platelet-endothelium adhesion molecule gene *Itga4*, component of the soluble guanylyl cyclase (a nitric acid sensor) *Gucy1a2*, and platelet factor *Pf4*. Noticeably, multiple immune system genes were upregulated in both strains, including *Cd48, Cd52, Ly96, Ifi203, Mrc1, Pf4, Ctss and H2-Aa* (**Fig7d)**.

Analysis of plasma from both strains with autoantigen arrays revealed continued divergence in immune activity. APOE cKO mice largely upregulated antibody responses to autoantigens at week 4 of HFD, consistent with the known role of APOE repressing immune activation ^18,25,27^. In contrast, D374Y displayed significant down-regulation of multiple autoreactive IgM, whereas only 3 IgG responses were affected (**Fig7e)**.

Finally, we next targeted CD8+ T cells to determine if a component of the immune system could influence this very early stage of atherosclerosis initiation. We confirmed that anti-CD8a antibodies successfully depleted CD8+ T cells, as previously reported ^28^ (**Fig8a)**. CD8+ T cell depleted APOE cKO mice selectively downregulated 6 genes in the aorta compared to APOE cKO with an intact immune system, as determined by mRNA-sequencing (**Fig8b)**. Only one gene was upregulated in APOE cKO mice, the T cell derived elastase *Serpina3n*, and this gene was the only gene differentially regulated in D374Y mice subjected to CD8a depletion (**Fig8c)**. In summary, mice subjected to continuous dyslipidemia begin to show convergent molecular responses in the aorta, and these changes can be modulated by T cells.

**Figure 8.**
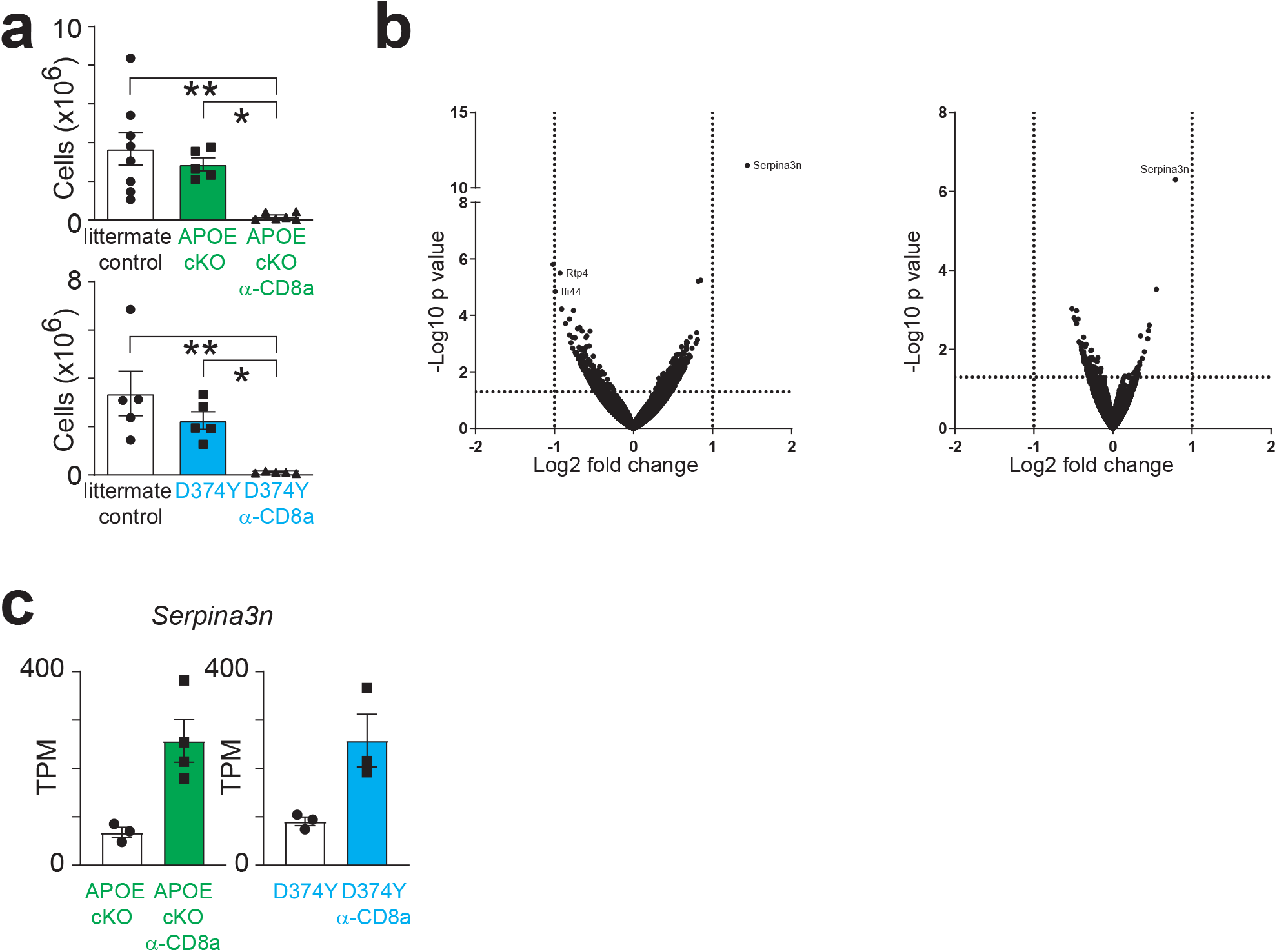
Response of the vasculature to loss of CD8+ T cells in inducible dyslipidemic mice. **A**. Absolute cell numbers of CD3+CD8+ cells from mice of the indicated genotypes and mice treated with anti-CD8a depleting antibodies as determined by flow cytometric analysis of the spleen. All mice are female and were studied at 4 weeks after tamoxifen dosing and start of high-fat diet. **B**. Volcano plot for gene expression differences in APOE cKO injected with anti-CD8a depleting antibodies (4 RNA-seq experiments) compared to PBS (3 RNA-seq experiments) and D374Y mice with anti-CD8a depleting antibodies (3 RNA-seq experiments) compared to PBS (3 RNA-seq experiments). **C**. Upregulation of *Serpina3n* in both APOE cKO and D374Y mice in responses to CD8+ T cell depletion. Plots are ± SEM and each dot corresponds to one mouse. Statistical analysis was performed with one-way ANOVA. *p < 0.05, **p < 0.01 compared to control mice.

## Discussion

We have investigated how a transition to dyslipidemia can initiate atherosclerosis in adult mice. To do so, we engineered strains that can acutely convert to an atherogenic lipid profile and observed microscopic accumulation of APOB at anatomical sites of plaque predilection. We have therefore captured the very first steps in the genesis of an atherosclerotic plaque. Notably, this is not possible in adult mice with current legacy strains, as we observed prominent APOB accumulation already at week 10 in *Apoe*^−/−^ and *Ldlr*^−/−^ strains, and aortic gene expression changes can be detected at week 6 in *Apoe*^−/−^ mice ^29^. Our observations in our new inducible models are consistent with the evidence based fact that raising plasma levels of apoB-lipoproteins can act as a rheostat for atherosclerosis development; whereby the higher the concentration the more rapid the lesion formation ^30^.

In response to dyslipidemia, we observed rapid APOB accumulation in the atherosclerosis prone lesser curvature intima of aortic arch. The same area was populated by CD68+ cells in both APOE cKO and D374Y mice, as well as their littermate controls. This is in line with previous studies showing that CD68+ cells are present in healthy mouse arteries, especially in the lesser curvature of the aortic arch, and the numbers increase in early atherosclerosis ^31,32^. Compared to littermate controls, there was an increase in CD68+ cell size in the D374Y strain, potentially indicating activation and lipid accumulation in the resident myeloid cells. It is a possibility that this activation could be the starting point of recruitment of other inflammatory cells to the nascent lesion. Future studies using spatial transcriptomics and single-cell approaches will be useful in identifying the individual cell types at atherosclerosis onset.

The mRNAseq experiments revealed diverse aortic gene expression changes in the two models. In APOE cKO at day 7 or day 10, the differentially regulated genes involved many that are related to stress response (*Egr1*, *Rnh1*, and *Creb3l2*). *Egr-1*, early growth response 1, is known regulatory factor in vascular injury, and its expression is increased in atherosclerotic lesions ^33^. It represses vascular repair mechanisms, for example by supressing transforming growth factor beta expression ^34^. In line with this, *Tgfb2*, which has a role in epithelial-mesenchymal transformation ^35^, was downregulated at day 10 and TGFB1 protein was decreased in the plasma of APOE cKO strain. Many genes related to extracellular matrix, coagulation or healing after vascular injury were also differentially regulated (*Fgb*, *Kng1*, *Serpine-1*). *Serpine-1* encodes protein plasminogen activator inhibitor-1, which inhibits cell migration ^36^ and its loss-of-function variant is associated with high cardiovascular risk ^37^. Additionally, several smooth muscle cell and vascular function related genes were downregulated. *S1pr1*, sphingosine-1-phosphate receptor 1 may indicate impairment of vascular relaxation ^38^, transcription factor *Klf15* has also been observed in human atherosclerosis resulting in changes in vascular cell adhesion molecule-1 (VCAM-1) as well as vascular endothelial growth factor (VEGF) signalling ^39^ and the VEGF-signalling related genes *Kdr* (encoding VEGFR2) and *Peli1* were also differentially regulated. In conclusion, the early responses to elevated plasma cholesterol in APOE cKO mice are related to vascular cell biology, and emphasize a specific role for APOE in vascular function ^40^.

The gene expression changes in D374Y mice diverged from the ones seen in APOE cKO at early time points. Induction of dyslipidemia in D374Y mice led to increased expression of several genes involved in fatty acid metabolism, especially the acyl-coenzyme A (CoA) reactions (*Acsm3*, *Acly*, *Acsl1*, *Acsl5*) and other fatty acid and triglyceride metabolism *(Dlat*, *Dgat1*, *Ehhadh*, *Lpl*, *Scd1*, *Dbi*, *Fabp4*). These changes might be a response to the elevation in the plasma triglycerides specific for the D374Y strain. *Acly*, ATP-citrate lyase, is an enzyme linking glucose catabolism and lipogenesis. Importantly, it catalyzes production of the high-energy biosynthetic precursor acetyl-CoA, a fuel used in the synthesis of cholesterol and fatty acids and now being considered a target for a novel lipid-lowering drug bempedoic acid ^41^. Several genes linked to cardiovascular disease were also upregulated, such as fatty acid binding protein 4 *Fabp4* ^42^, RAS, dexamethasone-induced 1 *Rasd1* ^43^, and complement component *C3* ^44^. However, most noticeable was the increased expression of factors linked to macrophage activation such as the bactericidal perforin *Mpeg1*, the prostaglandin E receptor 3 *Ptger3*, the low affinity immunoglobulin gamma Fc region receptor IV *Fcgr4*, and the microsomal glutathione S-transferase 1 protein *Mgst1*. Myeloid associated genes with an established critical functions in atherosclerosis development were also identified, including the scavenger receptor *Cd36* ^45^, the chemokine *Cxcl9* ^46^, lipid antigen presentation receptor *Cd1d1* and most importantly multiple genes of the major histocompatibility complex (MHC). Taken together, the gene expression changes in D374Y in response to early dyslipidemia reflect metabolic adaptation and immune activation, especially in the myeloid compartment.

The two mouse models also differed in their systemic inflammatory responses to dyslipidemia. Our results add further weight for a role of APOE in inhibiting systemic adaptive immune responses typified by GC formation and DC activation ^26,47–49^. However, we could not detect splenic GCs in our D374Y strain, consistent with our previous study with *Ldlr*^−/−^ mice ^18^. In line with this, the autoantibody responses were greater in APOE cKO compared to D374Y, the only exception being the IgG antibodies against modified LDL in D374Y. Our two strains will be helpful in future studies for discriminating whether physiological responses are really due to dyslipidemia or can instead be explained by loss of APOE.

Analysis of aortas after 4 weeks of HFD-enhanced dyslipidemia revealed a convergence in gene expression between the two models. Reassuringly, multiple immune genes were similarly upregulated in both strains compared to controls, including *Cd48, Cd52, Ly96, Ifi203, Mrc1, Pf4, Ctss and H2-Aa* as well as a number of extracellular matrix proteins. As atherosclerotic plaques in *Apoe*^−/−^ and *Ldlr*^−/−^ at the macroscopic level appear quite similar, this implies that although initiation responses are markedly different in these inducible strains, the pathological endpoint begins to coalesce around several overlapping processes.

Underlining the close relationship of the immune system and the vascular response to dyslipidemia, we studied how CD8a depletion affects the gene expression in the two models. In APOE cKO, the effects of CD8a depletion were greater than in D374Y, yet only a very few genes were differentially regulated. The only gene affected by CD8a depletion in D374Y was *Serpina3n*, which was upregulated also in APOE cKO. In addition to being an acute phase protein, serine protease inhibitor Serpina3n or its human orthologue SERPINA3 is associated with atherosclerosis and abdominal aortic aneurysm ^50–52^. As we have now created an experimental system that allow for manipulation of the initial physiological response to dyslipidemia, we believe our models will be an ideal test-bed for targeting additional immune cell types, for vaccination strategies and other therapeutic approaches aimed at preventing or reducing progression of atherosclerosis.

In summary, we have developed two complementary mouse models that allow for studying how dyslipidemia can initiate atherosclerosis plaque formation within the vasculature, a process that remains largely unexplored. Loss-of-function mutations in APOE and LDLR as well as gain-of-function mutations in PCSK9 are present in human familial hypercholesterolemia (FH) patients, who are on the far extreme of a dyslipidemic spectrum of those individuals who will develop atherosclerosis. Nevertheless, even in the context of highly-inbred laboratory mouse strains, vasculature and immune responses greatly differ in response to acute hypercholesterolemia, dependent on the means by which concentrations of APOB-containing lipoproteins have been raised. Given the much more complex nature of human atherosclerosis, of which mouse models of FH represent a very small fraction, it is plausible that clinical disease may also have multiple initiating responses. Our studies also further underline that atherosclerosis has an intimate relationship with the immune system, even at the earliest phases of disease development.

## Acknowledgements

We thank Maria Ozsvar Kozma for expert help in performing ELISA measurements, Indu Raman for help with microarray, and Linda Haglund and Anneli Olsson for technical help in the laboratory.

This work was supported by the European Community’s Seventh Framework Program FP7-2007–2013 under Grant HEALTH-F2-2013-602114 (Athero-B-Cell), The Novo Nordisk postdoctoral fellowship programme at Karolinska Institutet, The Swedish heart lung foundation (Hjärtlungfonden – project grants 20190655 & 20190357) the Nanna Svartz Foundation, Finnish Foundation for Cardiovascular Research (Sydäntutkimussäätiö) and the Karolinska Institutet.

## Author contributions

SM initiated and designed the study. SH and SM carried out the animal experiments and flow cytometry. SH performed immunofluorescence, RNA isolation and plasma lipid measurements. VS and SM performed bioinformatics. JM and MJ measured plasma lipids and lipoprotein profiles. CB and VM performed ELISA. GC and QZL performed autoantigen array. SH and SM analyzed the data and wrote the manuscript with input from all the authors.

## Data availability

The RNA-seq data reported in this study are available in GEO under accession number xxx

## Abbreviations

apoB: Apolipoprotein B
APOE: Apolipoprotein E
APOE cKO: Conditional knockout of APOE
CoA: Coenzyme A
D374Y: Conditional knock-in of hPCSK9 D374Y
DAPI: 4′,6-diamidino-2-phenylindole
DC: Dendritic cell
ELISA: Enzyme-linked immunosorbent assay
FDR: False discovery rate
FH: Familial hypercholesterolemia
GC: Germinal center
HFD: High fat diet
hPCSK9 D374Y: human gain-of-function variant of PCSK9
IgG: Immunoglobulin G
IgM: Immunoglobulin M
LDL: Low-density lipoprotein
LDLR: Low-density lipoprotein receptor
MDA-LDL: Malondialdehyde modified LDL
MHCII: Major histocompatibility complex II
mRNAseq: messenger RNA sequencing
PCSK9: Proprotein convertase subtilisin/kexin type 9
PDGFB: Platelet-derived growth factor subunit B
RPM: Mapped reads
TGFB: Transforming growth factor beta
TPM: Transcripts per million reads mapped
VCAM-1: Vascular cell adhesion molecule-1
VEGF: Vascular endothelial growth factor
VLDL: Very low-density lipoprotein

## Materials and Methods

### Mice and experimental diets

The APOE cKO mice have been described earlier ^18^ and were maintained on the C57BL/6 genetic background. All mice were housed in specific pathogen free vivarium at Karolinska institutet. Two to eight mice were housed in each cage. The light/dark period was 12h/12h, and all the mice had *ad libitum* access to food and water. Mice in breeding were fed chow diet R36 (12.6 MJ/kg, 18% protein, 4% fat; Lantmännen, Sweden). Experimental mice received chow diet (R70, Lantmännen, Sweden, 12.5 MJ/kg, 14% protein, 4.5% fat) or HFD (R638, Lantmännen, Sweden, 15.6 MJ/kg, 17.2% protein, 21% fat, 0.15% cholesterol) as stated in each experiment. Stockholm Board for Animal Ethics approved the experimental protocols.

### Generation of hPCSK9 D374Y mouse strain

We created a conditionally-activated hPCSK9 D374Y gain-of-function mouse model by inserting D374Y mutated hPCSK9 into *Rosa26* locus. The ROSA26 gene-targeting vector was constructed from genomic C57BL/6N mouse strain DNA (Genoway, Lyon, France). PCSK9 D374Y human sequence was inserted downstream of a lox-STOP-lox cassette. When the floxed STOP cassette is removed by CRE recombinase, human PCSK9 D374Y expression is driven by the CAG promoter. The linearized targeting vector was transfected into C57BL/6 embryonic stem cells (ES cells) (genOway, Lyon, France) according to genOway’s electroporation procedures. G-418 resistant ES cell clones were subsequently validated by PCR, using primers hybridizing within and outside the targeted locus, and Southern blot, to assess the proper recombination event on both side of the targeted locus. Recombined ES cell clones were microinjected into C57BL/6 blastocysts, and gave rise to male chimeras with a significant ES cell contribution. Breeding were established with C57BL/6 mice to produce the heterozygous inducible human PCSK9 D374Y line. These *ROSA2*^*PCSK9D374Y*^ mice were crossed with *ROSA26*^*CreERt2*^ mice, creating *ROSA26*^*CreERt2/PCSK9D374Y*^ experimental mice. Littermates without the D374Y insert (*Rosa26^CreERt2/+^ or Rosa26 ^CreERt2/ CreERt2^*) were always used as controls.

### Induction of dyslipidemia

Mice of both sexes were used in the studies and they were induced at the age of 10-14 weeks unless otherwise specified. Induction of the models with oral dose of tamoxifen was performed as we previously described^18^. We administered 9 mg of tamoxifen dissolved in 150μl peanut oil + 10% ethanol, via single-dose oral gavage, to experimental mice and their littermate controls. The induction efficiency was evaluated by measuring the plasma cholesterol levels. Occasional APOE cKO and D374Y mice that did not show the expected elevation in cholesterol levels were excluded from the study.

### Plasma lipid analyses

For the characterization of the cholesterol kinetics in the D374Y strain, tail vein blood samples were taken before tamoxifen dosing as well as at three days after it. Blood was drawn to EDTA-coated tubes and centrifuged at room temperature for 12 minutes at 2500 G. Plasma was separated and stored at −80°C for further analyses. For other plasma lipid analyses, mice were euthanized by carbon dioxide asphyxiation. Blood was drawn via cardiac puncture into EDTA-coated tubes, centrifuged, and stored as described above. The plasma total cholesterol and triglycerides were measured from plasma with enzymatic colorimetric kits (Randox laboratories) according to manufacturer’s instructions. Phospholipids (PL) were measured with Phospholipids C kit (Fujifilm, Wako Diagnostics). Plasma free (non-esterified) fatty acid concentrations were measured by an enzymatic colorimetric method (NEFA-HR(2), Wako Chemicals). Plasma concentration of glycerol was determined by an enzymatic colorimetric assay (Free glycerol FS, DiaSys, Diagnostic Systems GmbH).

Lipoprotein fractionation was performed for plasma pools of four to six mice by fast performance liquid chromatography method ^53^. The concentration of cholesterol, triglycerides and PL in each fraction was measured by enzymatic methods using CHOD-PAP kit (Roche Diagnostics) GPO-PAP kit (Roche Diagnostics) and Phospholipids C kit (Fujifilm, Wako Diagnostics), respectively.

### Histology and immunofluorescence

The mice were euthanized with carbon dioxide and perfused by infusing PBS via left ventricle. The aortic arch was dissected carefully under dissection microscope and mounted to optimal cutting temperature (OCT) compound for the analysis of histology. Each aorta was longitudinally sectioned at −20°C. Serial 10 μm thick sections were thaw-mounted to slides which were subsequently fixed with ice-cold acetone for 10 minutes and stored at −20°C. The presence of APOB containing lipoproteins and CD68+ cells was analysed by immunofluorescent staining. The slides were thawed and sections were incubated in washing buffer (1x Tris-buffered saline with 0.1 % Tween20) for five minutes. Blocking was performed with Avidin-biotin blocking kit (Vector) according to supplier instructions, and with 5% BSA for 30 minutes. Primary antibodies against CD68 (clone MCA1957, Serotec, 1:10000) and APOB (clone 20-AG40, Fitzgerald, 1:10000) were incubated at +4°C over night. The secondary antibody (biotinylated anti-goat IgG (Vector), 1:300) was incubated for 1 hour at RT followed by 1 h incubation with streptavidin-DyLight 647 (Vector, 1:300) and anti-rat DyLight 594 (Abcam, 1:300) and subsequently 4′,6-diamidino-2-phenylindole (DAPI) (Invitrogen, 1:50000) for 20 minutes. The sections were washed 3 × 3 minutes in washing buffer after each incubation step. The slides were mounted with fluorescent mounting media (Dako). Negative control stainings were performed by omitting the primary antibodies from the protocol. For CD68+ cells, staining in spleen was used as a positive control. Immunofluorescence images were taken with Nikon Ti-2E confocal microscope using NIS Elements software. Images with 10x magnification were taken from the ascending aorta, between the aortic root and innominate artery branch, including lesser and greater curvature. Two to three aortic sections per mouse were analysed in FIJI.^54^. APOB staining was quantitated in the lesser curvature of ascending aorta, by defining a segmented line (width 5 pixels, 6.2 μm, length 1mm) in the intima. The mean APOB staining intensities along the segmented line were calculated for each mouse and for each strain. Additionally, the average number and size of CD68+ cells in lesser curvature was analysed. The images of CD68 staining channel were all thresholded with fixed pre-defined parameters to detect CD68+ staining, and the CD68+ cells attached to the inner curve intima were counted (number of cells / mm intima and average cell size, μm^2^) by particle analysis plugin in FIJI.

### Messenger RNA sequencing

All mRNA sequencing was performed on female mice. The mice were euthanized with carbon dioxide and perfused by infusing PBS via left ventricle. The thoracic aorta, from ascending aorta to the level of diaphragm, was carefully dissected under a dissection microscope and all visible adventitia, perivascular fat and surrounding tissue was removed paying attention to avoid thymus contamination. Aortas were stored in RNALater (Qiagen) at −20°C until RNA extraction. The aortas were solubilised in Qiazol lysis reagent using Tissuelyser (Qiagen) and extracted RNA was isolated to upper fraction by chloroform. Purification was performed with RNeasy mini kit including on-column DNAse treatment, using a Qiacube robot, according to manufacturer’s instructions. RNA integrity was evaluated with Agilent 2100 Bioanalyzer (Agilent Technologies). The RNA integrity numbers exceeded 8.5 in all samples. RNA was selected using Poly(A) RNA Selection Kit (Lexogen) and sequencing libraries prepared with Lexogen QuantSeq V2. DNA fragments 200-800 bp for RNA-seq were selected. Cluster generation and sequencing was carried out by using the Illumina HiSeq 2500 system with a read length of 50 nucleotides (single-read) and aligned to the mouse transcriptome (genome assembly version of July 2007 NCBI37/mm9) using TopHat version 1.4.1 ^55^. Reads per gene was counted using HTseq version 0.5.3 ^56^ with the overlap resolution mode set to union. Reads were visualised using the IgB browser. Analysis of differential expression of mRNA was using the DeSeq2 software at default settings.

### Detection of cell types by flow cytometry

Spleens were harvested and prepared to single cell suspensions by pressing through sterile 100 μm mesh size cell strainers. Erythrocytes were lysed for 3 minutes with 3 ml of EL buffer (Qiagen). After Fc blocking for 20 minutes, the cells were stained with conjugated antibodies on ice for 30 minutes. The following antibody clones were used: CD3ε (PB 500A2 or PerCP 145-2C11), CD4 (APCH7 GK1.5), CD8 (FITC 53-6.7), CD44 (PE IM7), CD62L (APC MEL-14), CD25 (PE-Cy7 PC61) B220 (APC-Cy7 RA3-6B2), CD19 (APC 1D3), GL7 (APC GL-7), CD95 (PE-Cy7 Jo2), IgD (PerCP 11-26c.2a), CD11c (APC HL3), CD64 (BV421 x54-5/7.1), CD172a (BV711 P84), XCR1 (PE ZET), MHCII (AF 700 M5/114.15.2).Immune cell populations were defined as: CD4 T-cell (CD3+CD4+CD8−), CD8 T-cell (CD3+CD4−CD8+), naïve CD4 T-cell (CD3+CD4+CD8−CD44−CD62L+), effector CD4 T-cell (CD3+CD4+CD8−CD44+CD62L−), B-1 (CD19+B220lo), B-2 (CD19+B220+), germinal center B (CD19+B220+IgD-GL7+CD95+), cDC1 (CD3−CD19−CD64−CD11c+XCR1+CD172a−) and cDC2 (CD3−CD19−CD64−CD11c+XCR1-CD172a+). The samples were acquired with FACSVerse (BD Biosciences) flow cytometer or Cytek Northern Light 3000 spectral flow cytometer (Cytek) and analyzed with FlowJo software.

### ELISA

Concentration of hPCSK9 was measured from the D374Y and littermate control mouse plasma obtained from tail vein blood before tamoxifen administration as well as at three days after it, and from cardiac puncture blood derived plasma at 10 days after tamoxifen dosing. The measurement was performed by Enzyme-Linked Immunosorbent Assay (ELISA) using the Human PCSK9 Quantikine ELISA kit (R&D Systems, Catalog no. DPC900, Minneapolis, USA) according to kit instructions. The lower limit of quantification (LLOQ) of hPCSK9 in mouse plasma was 625 pg/ml.

The presence of antibodies against modified LDL was evaluated by ELISA. Modified LDLs (copper oxidized and malondialdehyde-modified) were prepared according to previously described methods.^57^ For detection, alkaline phosphatase-labeled goat antibodies against mouse IgM μ chain (Sigma-Aldrich) and mouse IgG1 γ chain (Sigma-Aldrich) were used. T15/E06-clonospecific antibodies were detected with a chemiluminescent assay as described previously.^18^

### Plasma protein analysis by proximity extension assay

Concentrations of 88 different plasma proteins were evaluated by proximity extension assay (Mouse exploratory panel, Olink Proteomics).

### Autoantigen array

Plasma IgM and IgG antibody reactivities to a selection of 120 autoantigens were measured by Microarray core facility of UT Southwestern Medical Centre using a protein array method. ^58^ Briefly, plasma samples were diluted 1:100 and incubated with arrays containing 120 antigens and control proteins. IgM and IgG antibodies binding with the antigens were measured with Cy5-conjugated anti-mouse IgM and Cy3-conjugated anti-mouse IgG (Jackson ImmunoResearch Lab). A score for each antibody (Ab score) was calculated based on the net fluorescent intensity (NFI) and signal-to-noise ratio (SNR) using the formula: Ab score = log_2_ (NFI*SNR+1).

### CD8 depletion

CD8 cells were depleted with anti-CD8a antibody (Rat IgG2b anti-mouse CD8α YTS 169.4, BioXcell). D374Y and APOE cKO mice and their littermate controls were injected 250 μg antibody in 150 μl saline or similar volume saline intraperitoneally once weekly. Five days after the first injections, tamoxifen was dosed via oral gavage and the mice were switched to receive HFD. The diet and weekly injections continued for four weeks.

### Statistical analysis

Data are presented as means ± SEM Unpaired Student’s t test used for comparing two experimental groups as indicated. One or Two-way ANOVA followed by fisher’s LSD, or repeated measures ANOVA was used for comparisons of more than two groups as indicated. P < 0.05 was considered significant. Statistical analyses were performed using GraphPad Prism software.

---

**Supplementary Figure 1.**
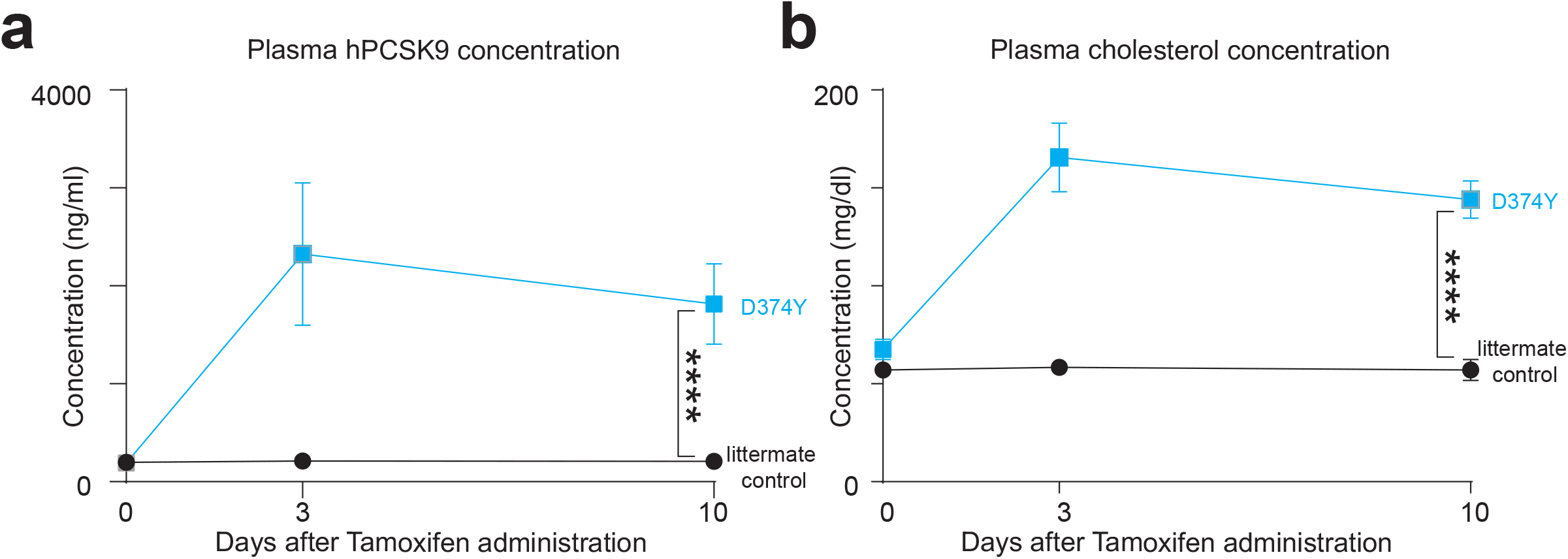
Plasma hPCSK9 and total cholesterol concentrations in D374Y. **A.** Enzyme-linked immunosorbent assay for plasma hPCSK9 protein in inducible D374Y mice (blue line) and their littermate controls (black line) before and at 3 and 10 days after tamoxifen dosing. n=2-6. **B.** Plasma total cholesterol measurements in the same time points as in A. n=4-6. All plots are ± SEM, n=3-. Statistical analysis was performed with mixed model 2-way ANOVA. ****p < 0.0001.

**Supplementary Figure 2.**
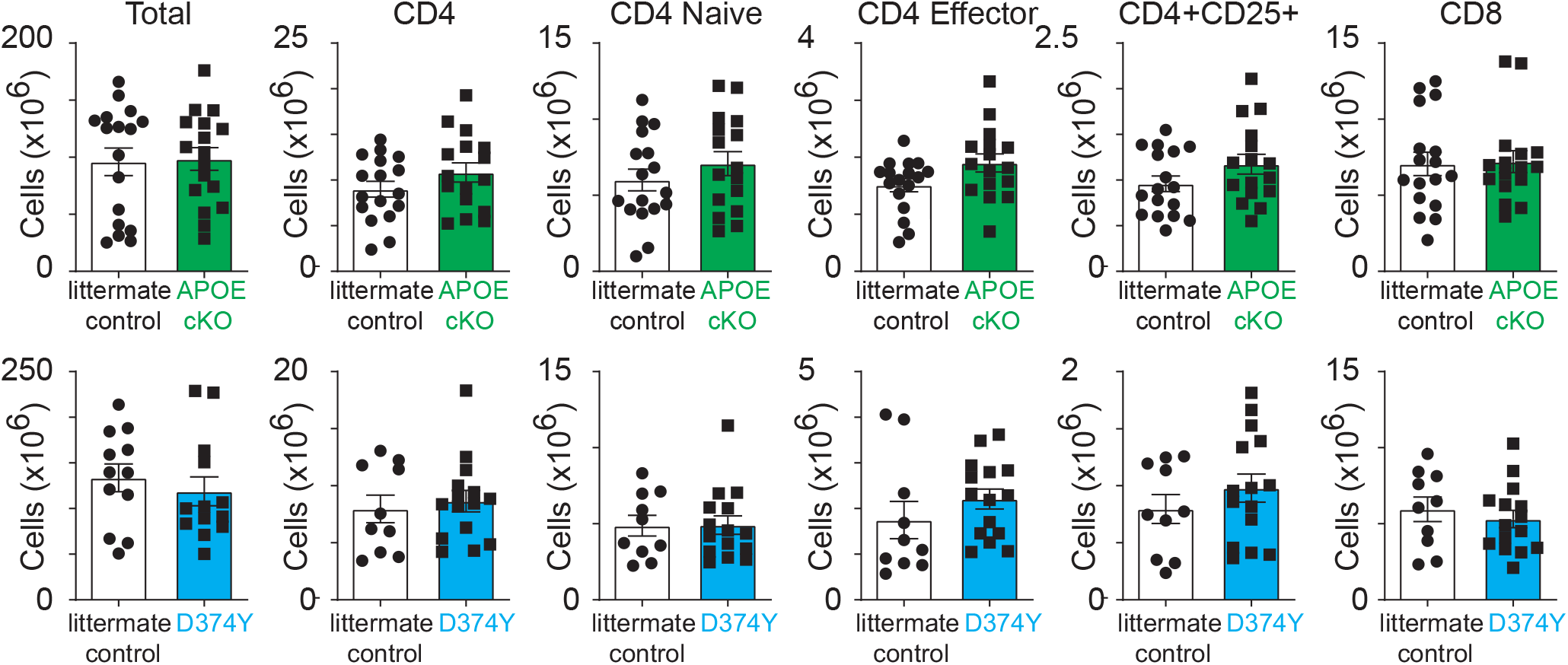
T cell populations in the spleen. Cell numbers of indicated T cell subsets as determined by flow cytometric analysis of the spleen from APOE cKO, D374Y and their respective littermate controls at day 10 following tamoxifen administration. T cells were defined as follows; CD4 T-cell (CD3+CD4+CD8−), CD8 T-cell (CD3+CD4-CD8+), naïve CD4 T-cell (CD3+CD4+CD8−CD44-CD62L+), effector CD4 T-cell (CD3+CD4+CD8−CD44+CD62L-) and CD4+ CD25+ (CD3+CD4+CD8−CD25+).

